# Senescence-induced cellular reprogramming drives cnidarian whole-body regeneration

**DOI:** 10.1101/2022.09.02.506310

**Authors:** Miguel Salinas-Saavedra, Febrimarsa, Gabriel Krasovec, Helen R Horkan, Andreas D Baxevanis, Uri Frank

**Affiliations:** Centre for Chromosome Biology, School of Biological and Chemical Sciences, University of Galway, Galway, Ireland; Computational and Statistical Genomics Branch, Division of Intramural Research, National Human Genome Research Institute, National Institutes of Health, Bethesda, MD 20892 USA

## Abstract

Cell fate stability is essential to maintaining ‘law and order’ in complex animals. However, high stability comes at the cost of reduced plasticity and, by extension, poor regenerative ability. This evolutionary trade-off has resulted in most modern animals being rather simple and regenerative or complex and non-regenerative. The mechanisms mediating cellular plasticity and allowing for regeneration remain unknown. We show that signals emitted by senescent cells can destabilize the differentiated state of neighboring somatic cells, reprogramming them into stem cells that are capable of driving whole-body regeneration in the cnidarian *Hydractinia symbiolongicarpus*. Pharmacological or genetic inhibition of senescence prevented reprogramming and regeneration. Conversely, induction of transient ectopic senescence in a regenerative context resulted in supernumerary stem cells and faster regeneration. We propose that senescence signaling is an ancient mechanism mediating cellular plasticity. Understanding the senescence environment that promotes cellular reprogramming could provide a new avenue to enhance regeneration.

## INTRODUCTION

Regenerative abilities of animals are irregularly scattered among taxa but are largely reverse-correlated with their structural complexity. With some exceptions, simple animals such as sponges, cnidarians, and planarians can regenerate whole-bodies from tissue fragments while complex animals such as vertebrates have more limited capabilities to restore lost body parts (Bely and Nyberg, 2010). Regeneration requires a certain level of growth plasticity that can be realized by a resident pool of stem cells or by mechanisms that induce reprogramming of differentiated cells to provide proliferative progenitors (Tanaka and Reddien, 2011). However, the high plasticity that allows for whole-body regeneration may also compromise the integrity of complex structures and increase malignancy risk (Sánchez Alvarado and Yamanaka, 2014; Slack, 2017). Therefore, high plasticity and structural complexity rarely co-occur in one species.

A major question arising from this line of arguments concerns the nature of the mechanisms that mediate cellular plasticity in regenerative animals. If regeneration was a primitive trait in metazoans (Bely and Nyberg, 2010), these mechanisms have been lost or became ineffective in non-regenerative taxa. Therefore, studying animals with high regenerative abilities could provide insight into the lack of regeneration in other animals. In the long term, this information might be harnessed to induce plasticity in complex, non-regenerative animals, enhancing their regenerative abilities under controlled conditions.

Studying regeneration in a cnidarian, we show that senescence signaling can drive somatic cell reprogramming in a regenerative context. Our results, combined with data from other animals, suggest that senescence is an ancient mechanism in animals, used to convey a stress signal to surrounding tissues, thereby driving a regenerative response. We propose that the ability to respond to a senescence signal is one of the factors underlying differential regenerative abilities in modern metazoans.

## RESULTS

### Stem cell-less tissues can regenerate whole animals that include stem cells

The cnidarian *Hydractinia symbiolongicarpus* — a relative of jellyfishes and corals — is a highly regenerative animal that is able to regrow a lost head within 3 days post amputation (dpa) (Bradshaw et al., 2015). *Hydractinia* head regeneration is driven by a population of adult migratory stem cells, known as i-cells (Gahan et al., 2016), that are normally restricted to the lower body column of the animal and can differentiate into both somatic cells and gametes (DuBuc et al., 2020). These i-cells, which can easily be visualized via *Piwi1* expression, migrate to the injury site post-amputation to restore the head. However, the heads of uninjured animals are devoid of i-cells (Figure 1A) (Bradshaw *et al*., 2015), so the head alone would not be expected to be able to regenerate a new body due to its lack of i-cells. Surprisingly, we have discovered that the amputated oral tips of heads (known as hypostomes) can indeed regenerate into a fully functional animal (Figure 1B) that contains i-cells despite having no i-cells immediately post-amputation (Figure 1C). We termed the i-cells that appeared *de novo* in regenerating hypostomes ‘secondary i-cells’, as opposed to primary i-cells that are generated during embryogenesis.

**Figure 1.**
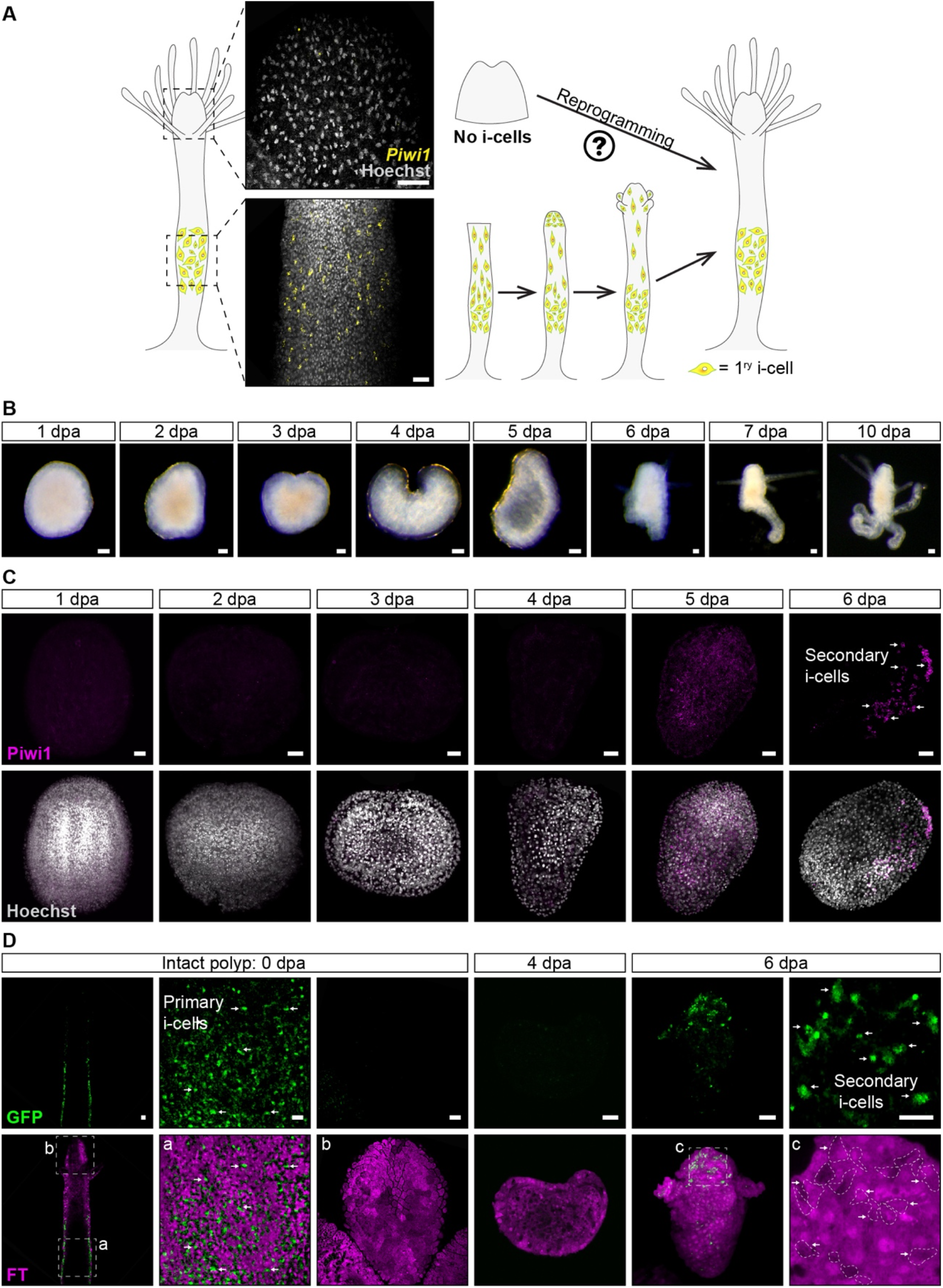
Secondary i-cells emerge in amputated hypostomes 6 dpa. **(A)** *Piwi1* mRNA SABER-FISH of an intact *Hydractinia* polyp. Primary i-cells are restricted to the lower part of the body but absent from the hypostome. **(B)** Amputated hypostomes can regenerate a fully functional animal within 6-10 dpa. **(C)** Piwi1 antibodies detect secondary i-cells in hypostomes, 6 dpa (arrows). **(D)** *In vivo* imaging of primary and secondary i-cells (arrows) in a *Piwi1*::FastFT reporter animal. In the intact polyp, primary i-cells (GFP^+^) do not overlap with the FT red fluorescence and are absent from the hypostome. However, in amputated hypostomes, secondary i-cells emerging at 6 dpa (depicted with arrows and dashed lines) overlap with the FT red fluorescence (c) observed at 4 dpa. Tissues do not express GFP at 4 dpa. This shows the reprogramming od somatic cells to secondary i-cells. Scale bars: 20 μm

To visualize the process of secondary i-cell appearance *in vivo*, we generated a *Piwi1* transgenic reporter animal that expressed a timer mCherry protein (Subach et al., 2009) and membrane GFP under the control of the *Piwi1* genomic control elements (Bradshaw *et al*., 2015). While the immature timer protein could not be observed under our microscopes due to the lack of a commercial filter, i-cells expressing a bright green membrane GFP could be viewed in the live translucent animal, while mature red timer protein was readily visible in differentiated cells (Figure 1D). The reporter transgene is silenced during differentiation, resulting in the early progeny of i-cells becoming red due to maturation of the timer protein and dim green due to the GFP’s half-life (Figure 1D). Hypostomes were then isolated from transgenic *Piwi1* reporter animals and examined under a fluorescence microscope to exclude the presence of any bright green i-cells; these hypostomes were then incubated in seawater (Figure 1D). Consistent with the experiments described above, GFP^+^ i-cells reappeared, *de novo*, at 6 dpa in an otherwise mature red mCherry^+^ background (Figure 1D). Hence, we concluded that, following amputation, a yet unknown mechanism induces the reprogramming of differentiated somatic cells back to *Piwi1*^+^ i-cells that then drive whole body regeneration.

Intact *Hydractinia* hypostomes lack i-cells and cycling cells (Bradshaw *et al*., 2015). Based on EdU incorporation, we established that a wave of DNA synthesis occurs between 3-4 dpa (Figure 2A). Secondary Piwi1^+^ i-cells appeared in the hypostome tissue around 6 dpa (Figure 1). These cells had residual EdU signals, showing that they were derived from cells that were in S-phase two days earlier (Figures 2B and S1A) Treating hypostomes with hydroxyurea to arrest cells in S-phase did not completely abolish the upregulation of Piwi1 in some cells (Figure 2C). However, the level of Piwi1 expression in these cells was low as compared to i-cells assessed using immunofluorescence. Moreover, hydroxyurea-treated hypostomes did not regenerate and subsequently died, showing that proliferation occurring before secondary i-cell appearance is essential for complete reprogramming of somatic cells to fully functional i-cells.

**Figure 2.**
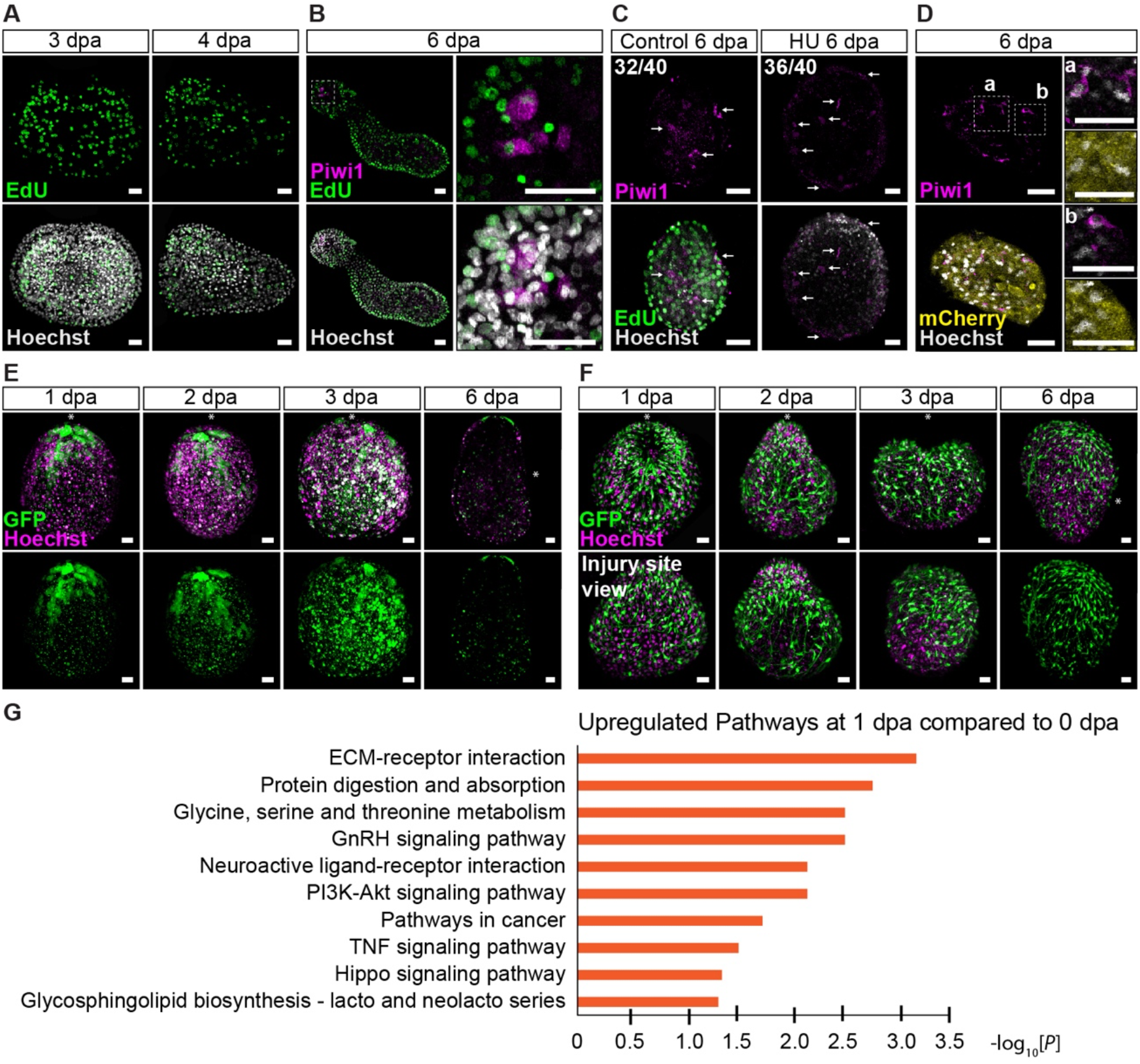
Cellular and molecular events accompanying whole-body regeneration from amputated hypostomes. **(A)** Cell cycle re-entry (EdU incorporation) takes place between 3 and 4 dpa. **(B)** Piwi1^+^ cells have residual EdU signals at 6 dpa after being incubated in EdU for 48 hrs between 3 and 4 dpa. **(C)** Few Piwi1^+^ cells appear in hydroxyurea-treated hypostomes but do cycle and are unable to contribute to whole-body regeneration. **(D)** Piwi1^+^ cells are also mCherry^+^, indicating the reprogramming of adult somatic cells. **(E)** *In vivo* imaging of amputated hypostomes from a *Wnt3*::GFP reporter animal 1, 2, 3, and 6 dpa. Lower panels show the GFP channel only. Asterisks indicate the oral pole pre-amputation. **(F)** *In vivo* imaging of amputated hypostomes from an *RFamide*::GFP reporter animal. Lower panels show the site of amputation. Asterisks indicate the oral pole pre-amputation. **(G)** Kegg Pathways upregulated in hypostomes, 1 dpa. Scale bars: 20 μm.

To verify that somatic cells undergo reprogramming to give rise to secondary i-cells, we amputated hypostomes from a *β-tubulin::FastFT* transgenic reporter animal. These animals express mCherry in all differentiated cells but not in i-cells (DuBuc *et al*., 2020). We allowed the hypostomes to develop secondary i-cells, then fixed and stained them with antibodies raised against Piwi1 and mCherry. We found cells that were double-positive, showing that cells that had expressed mCherry in the transgenic reporter animal (i.e., differentiated cells) reprogrammed into Piwi1^+^ i-cells (Figure 2D).

The *Hydractinia* main body axis is generated and maintained by Wnt/β-catenin signaling (Duffy et al., 2010; Plickert et al., 2006). We used a *Wnt3* transgenic reporter animal that expresses GFP in the oral-most tip of the hypostome (DuBuc *et al*., 2020) to dynamically follow the polarity of the isolated hypostomes. We found that GFP expression in the reporter animal lost its oral focus within 2-3 dpa, spreading around the tissue (Figures 2E and S1B). We repeated the experiment with an *Rfamide* transgenic reporter animal that expresses GFP in a stereotypic fashion in the nervous system of the head (Gahan *et al*., 2016). As with the *Wnt3* reporter, within 2-3 dpa, the GFP^+^ neurons lost their typical oral orientation and became disorganized (Figure 2F and S1B). We concluded that axis polarity is lost in isolated hypostomes within 3-4 dpa. Following secondary i-cell appearance, the hypostomes elongated and regained polarity, as visualized by *Wnt3* and *Rfamide* expression in transgenic reporter animals (Figures 2E-F; 6 dpa), developing into intact yet small animals.

### Senescence precedes regenerative reprogramming

We then aimed to identify the nature of the signal that induces the appearance of secondary i-cells in isolated hypostomes that were initially devoid of i-cells. We hypothesized that this signal is present in regenerating tissues a few days before both reprogramming and the emergence of secondary i-cells. Therefore, we extracted RNA from amputated hypostomes 0, 1, 3, and 6 dpa and sequenced their transcriptomes. Comparing the gene expression of each stage to its preceding one, we performed KEGG pathway analyses (Kanehisa et al., 2021) and identified cellular processes consistent with a senescence episode in 1 dpa hypostomes (Figure 2G; Supplemental File 1).

Senescence is a form of irreversible cell cycle arrest in animals, induced by stress, damage, oncogene expression, or telomere attrition (Rhinn et al., 2019; Roy et al., 2020). Senescent cells secrete a cocktail of factors, collectively known as the senescence-associated secretory phenotype (SASP) (Krtolica et al., 2001); these factors induce inflammation and enhance senescence and malignancy in neighboring cells (Krtolica *et al*., 2001; Paramos-de-Carvalho et al., 2021). Long-term accumulation of senescent cells in tissues is thought to contribute to organismal aging (Baker et al., 2011). Conversely, a role for senescence (and short-term senescence in particular) in cellular plasticity has been reported in mammals and other vertebrates (Da Silva-Alvarez et al., 2020; Paramos-de-Carvalho *et al*., 2021; Ritschka et al., 2017; Walters and Yun, 2020). However, its natural context of action is not well-understood. We hypothesized that amputation injury induces senescence in some cells that then emit a signal (through SASP) that promotes reprogramming of neighboring somatic cells. This would, in turn, produce stem cells that could initiate a regenerative process.

Following this line of reasoning, we looked for indicators of senescence in regenerating hypostomes. Cell cycle regulators such as p21 (encoded by *CDKN1A*) and p16 (encoded by *CDKN2A*) are known senescence markers in mammals (Hernandez-Segura et al., 2018). The *Hydractinia* genome encodes three *CDKN1A*-like genes (Figure S2) but no *CDKN2A*, the latter being vertebrate-specific. Our phylogeny could not resolve orthology between cnidarian and bilaterian CDKN1A proteins, suggesting that the *Hydractinia* genes were paralogs. Therefore, we called the *Hydractinia* three *CDKN1A*-like genes *Cyclin-dependent kinase inhibitor 1* (*Cdki1*), *Cdki2*, and *Cdki3*, respectively. Single-molecule fluorescence mRNA *in situ* hybridization showed that the three genes were expressed in some cells along the body column (Figures 3A and S3A) but only *Cdki1* was upregulated at the cut side of isolated hypostomes around 1 dpa (Figure 3B), being nearly undetectable in intact hypostomes (Figure 3A). Expression patterns of *Cdki2* and *Cdki3* did not visibly change following injury. An additional senescence indicator is senescence-associated β-galactosidase (SAβ-Gal) activity (Hernandez-Segura *et al*., 2018).

**Figure 3.**
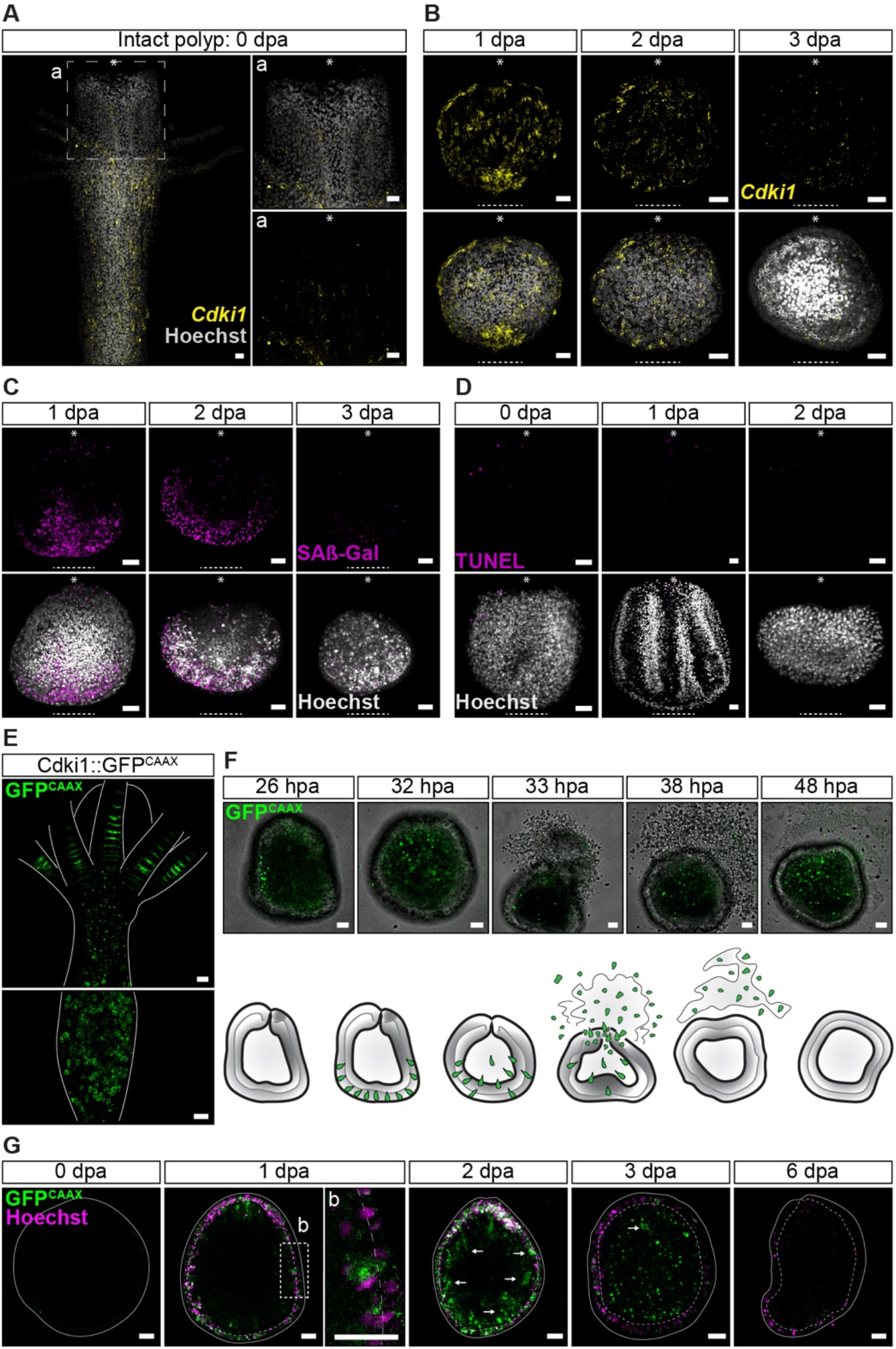
Senescence markers are transiently upregulated 1 dpa. **(A)** *Cdki1* mRNA SABER-FISH of an intact *Hydractinia* polyp. Low levels of *Cdki1* mRNA are present in the hypostome. **(B)** *Cdki1* mRNA SABER-FISH of amputated hypostomes 1, 2, and 3 dpa. *Cdki1* mRNA is upregulated at the site of injury (dashed line) at 1 dpa and dissipates 3 dpa. Asterisks indicate the oral pole pre-amputation. **(C)** SPiDER-βGal staining of amputated hypostomes 1, 2, and 3 dpa. βGal activity resembles *Cdki1* mRNA expression pattern at the site of injury (dashed line). Asterisks indicate the oral pole pre-amputation. **(D) γ**H2AX, H3K9me (arrows indicate the gastrodermis), and absence of apoptosis (TUNEL) at the site of injury (dashed line). Asterisks indicate the oral pole pre-amputation. **(E)** *In vivo* imaging of an intact *Cdki1*::GFP^CAAX^ transgenic reporter animal. **(F)** Time-lapse imaging (extracted from Supplemental Movie 1) of an amputated *Cdki1*::GFP^CAAX^ reporter hypostome at 26, 32, 33, 38, and 48 hpa. Interpretation of the images are shown as cartoon. **(G)** Time-lapse imaging of amputated hypostomes from a *Cdki1*::GFP^CAAX^ reporter animal showing the migration of senescent cells from the epidermis to the gastrodermis (arrows). No *Cdki1*::GFP^CAAX^ cells are present 6 dpa. Scale bars: 20 μm

Using SPiDER-βGal, a cell permeable fluorescent β-galactosidase probe, we found high SAβ-Gal activity around the injury site of hypostomes, concomitantly with *Cdki1* expression (Figures 3C and S4A). The above signals reached a peak around 1 dpa and vanished 2-3 dpa, indicating a transient, short-term senescence episode at the injury site post amputation.

In the closely related cnidarian *Hydra*, it has been shown that bisection induces apoptosis in i-cells, with these dying cells emitting a Wnt3 signal that drives proliferation and head regeneration (Chera et al., 2009). However, in *Hydractinia* regenerating hypostomes, no evidence for apoptosis was found using TUNEL staining (Figures 3D, S4B, and S4C), suggesting that senescence-induced regeneration in the absence of stem cells is distinct from apoptosis-induced repair where i-cells are present.

Given that senescent cells do not spontaneously die or normally exit senescence, the disappearance of senescence markers between the second and third dpa prompted us to investigate their fates. For this, we generated a *Cdki1* transgenic reporter animal that expressed membrane GFP^CAAX^ under the *Cdki1* genomic control elements. GFP^CAAX^ fluorescence in transgenic animals faithfully recapitulated *Cdki1* mRNA expression (Figure 3E), being upregulated at the injury site 1 dpa (Figures S3, S4B, and S4C). We amputated hypostomes from these animals and subjected them to *in vivo* time-lapse imaging (Figures 3F and S3D; Movie 1). Strikingly, these experiments showed that epidermal senescent cells translocate to the gastrodermal tissue (Figure 3G) and are then expelled, probably through the mouth, by regenerating hypostomes between 30-40 hours post amputation (hpa; Figures 3F-G, S3C, and S3D), consistent with the disappearance of senescence markers (Figure 3C). Therefore, whole-body regeneration from isolated hypostomes is accompanied by a short senescence period around 1 dpa, loss of tissue polarity within 3 dpa, a burst of proliferation around 3-4 dpa, and secondary i-cell appearance 6 dpa (Figure 4A). The process is completed by the re-establishment of tissue polarity, elongation, and morphogenesis.

**Figure 4.**
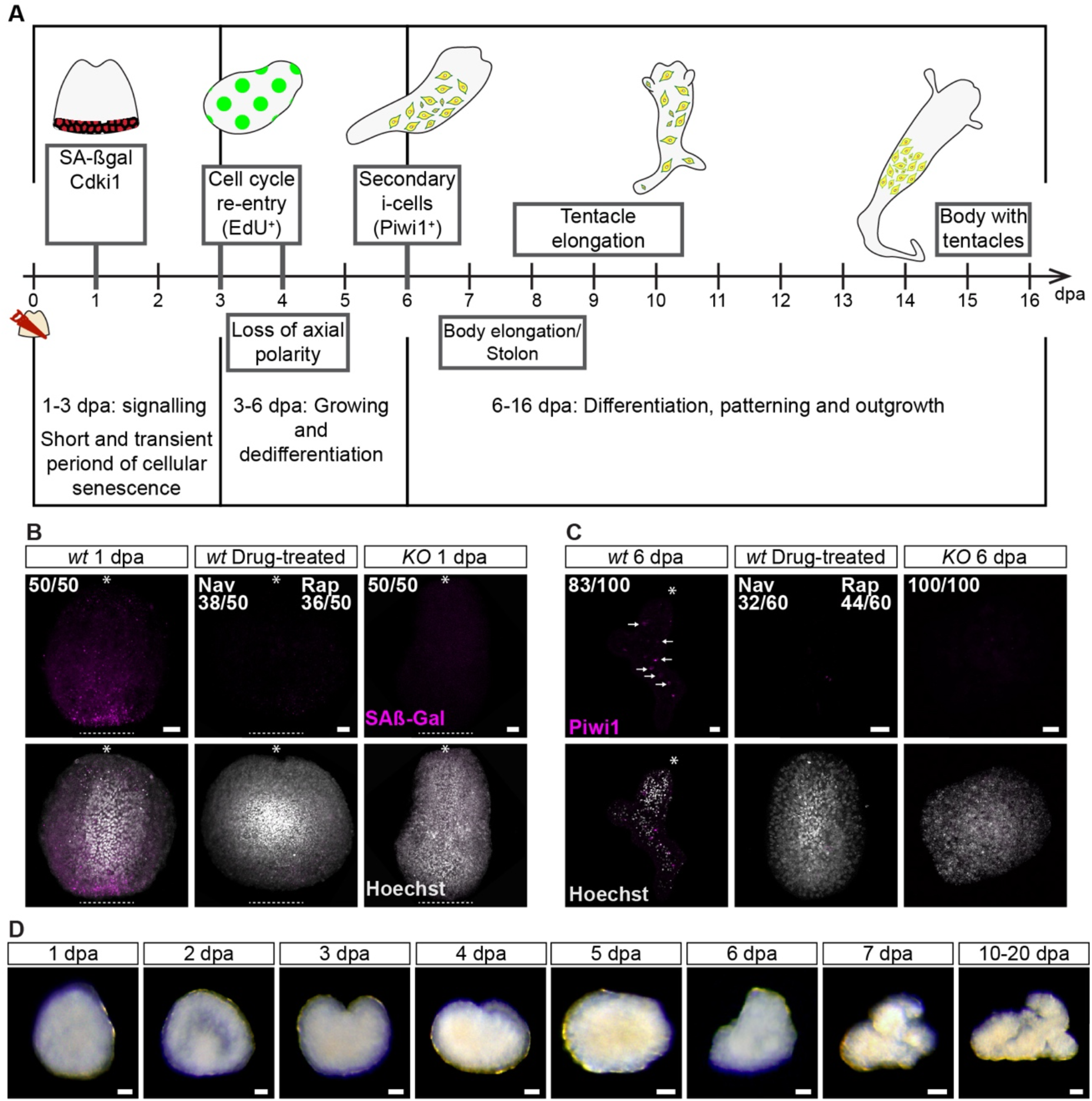
Transient senescence 1 dpa is required for somatic cell reprogramming. **(A)** Diagrammatic illustration of the major events accompanying whole-body regeneration from amputated hypostomes. **(B)** Hypostomes amputated from Navitoclax-treated and *Cdki1* KO animals do not develop cellular senescence at the site of injury 1 dpa (dashed line). Asterisks indicate the oral pole pre-amputation. **(C)** Hypostomes amputated from Navitoclax-treated and *Cdki1* KO animals do not develop Piwi1^+^ cells (arrows) at 6 dpa. Asterisks indicate the oral pole. **(D)** Amputated hypostomes from *Cdki1* KO animals do not regenerate within 20 dpa. Scale bars: 20 μm

### Senescence signaling is required and sufficient to induce reprogramming

To identify a functional link between the short period of senescence and the subsequent reprogramming of somatic cells, we used navitoclax, a senolytic drug (Chang et al., 2016), to inhibit senescence and study the effect of this manipulation on somatic cell reprogramming and secondary i-cell appearance. We exposed freshly amputated hypostomes to 1 μM navitoclax in seawater and followed them over 6 dpa. We found that navitoclax at this concentration inhibited senescence marker upregulation for the duration of treatment (Figure 4B). Furthermore, no secondary i-cells appeared in treated hypostomes at 6 dpa (Figure 4C). We repeated the experiments with rapamycin, an mTOR inhibitor, which yielded similar results.

To further address the requirement of senescence to reprogramming and exclude an unspecific effect of senolytic drugs, we used CRISPR-Cas9 to mutate the *Cdki1* gene (Figure S5A). Three short guide RNAs (sgRNAs) targeting the nuclear localization signal (NLS) and PCNA-interacting domain of *Cdki1* were designed and injected with recombinant Cas9 into fertilized eggs (Gahan et al., 2017). G0 embryos were allowed to develop into larvae, metamorphose, and grow to the young colony stage. They were then screened for mutations by genomic PCR and sequencing.

Confirmed mosaic mutants were grown to sexual maturity and crossed with wild type animals. Heterozygous G1 animals were identified by PCR and sequencing, grown to sexual maturity, and interbred to give rise to G2 homozygous knockout (KO) animals (Figure S5A). These animals developed normally to the larval stage, metamorphosed, and grew to apparently normal colonies that were able to regenerate amputated heads similar to those seen in wild type animals (Figure S5B). However, EdU analysis revealed that *Cdki1* KO animals had an abnormal, broader distribution of cycling cells, consistent with the loss of a cell cycle regulator (p<0.0001, n=20; Figures S5C-D; Supplemental File 2).

We amputated hypostomes from *Cdki1* KO animals and analyzed their behavior. We found that the absence of functional Cdki1 resulted in the loss of the senescence response at 1 dpa (Figure 4B) and absence of secondary i-cells at 6 dpa (Figure 4C). With the lack of i-cells, the KO hypostomes had not regenerated within 20 dpa (Figure 4D), remaining as amorphous tissue lumps that eventually died of starvation. Therefore, a short senescence signaling is essential for reprogramming.

Finally, we tested the ability of ectopic senescence to induce reprogramming in this specific context. To induce senescence, we employed an optogenetic approach using the Opto-SOS genetic cassette (Johnson et al., 2017). Cells carrying this construct respond to blue light by overactivation of the Ras pathway (Figure 5A), and overactivation of Ras is known to induce senescence (Serrano et al., 1997). We generated transgenic mosaic animals that carried the Opto-SOS construct, fused to mScarlet. Hypostomes were amputated and exposed to blue light for 12 hpa (Figure 5B) and we then observed the events accompanying their regeneration. As expected, we found that exposure to blue light induced enhanced SAβ-Gal activity (Figures 5C-D), and secondary i-cells appeared 5 dpa as opposed to 6 dpa in animals kept in the dark. At 6 dpa, the number of secondary i-cells was significantly enhanced in animals exposed to blue light comparing to those kept in the dark (p<0.0001, n=44; Figure 5E-F; Supplemental File 2). Taken together, a signal emitted by senescent cells that are transiently present following injury is essential and sufficient to induce the reprogramming of somatic cells to stemness in amputated hypostomes.

**Figure 5.**
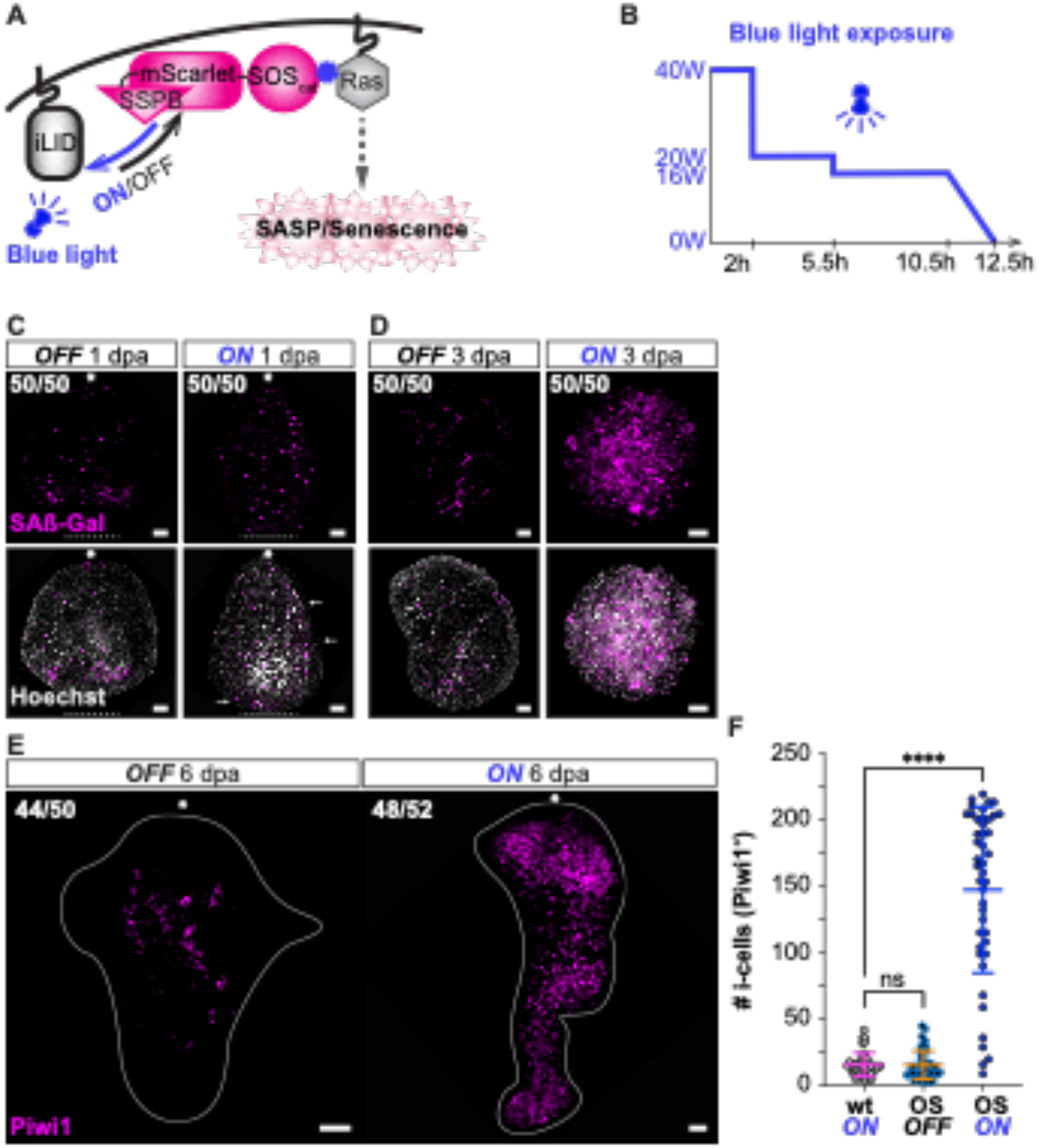
An optogenetic approach for ectopic induction of senescence during regeneration results in increased numbers of secondary i-cells. **(A)** A diagrammatic representation of the optogenetic activation of the Ras pathway when exposed to blue light. **(B)** Blue light exposure pattern during the experiments. **(C)** Ectopic senescence outside the site of injury at 1 dpa (dashed line) under blue light exposure. Asterisks indicate the oral pole pre-amputation. **(D)** Enhanced cellular senescence 3 dpa under blue light exposure. **(E)** Supernumerary secondary i-cells at 6 dpa under blue light exposure. **(F)** Quantification of the number of secondary i-cells at 6 dpa (p<0.0001, n=44). Amputated hypostomes were incubated during 1 dpa under three different conditions: Wild type under blue light exposure (wt ON), OptoSOS under darkness (OS OFF), OptoSOS under blue light exposure (OS ON). Scale bars: 20 μm.

## DISCUSSION

Animal somatic cells face contradictory requirements during the life history of individuals. On the one hand, cell fate stability is needed to maintain structural integrity and prevent malignancy while, on the other hand, stably differentiated cells prevent regeneration. Extant animals appear to deal with this problem using a spectrum of strategies. Morphologically simple animals such as cnidarians possess high cellular plasticity and regenerative powers. They can tolerate the presence of unstable cells because their morphological simplicity facilitates the shedding of unwanted cells (Figures 3F-G). By contrast, morphologically complex bilaterians such as mammals cannot afford to harbor developmentally plastic cells due to malignancy risk and the necessity to maintain complex structures. The results presented here are consistent with the notion that senescence signaling is one of the factors that mediate cellular plasticity (Rhinn *et al*., 2019). However, the ability to respond to a senescence signal has not been highly conserved across lineages during metazoan evolution.

A rudimentary response to senescence signals by increased plasticity is still present in modern mammals. This has been shown in mouse liver cells (Ritschka *et al*., 2017), skeletal muscles (Chiche et al., 2017), and by the discovery that a senescent environment facilitates reprogramming by OSKM factors (Mosteiro et al., 2016).

Finally, embryonic and induced pluripotent stem cells maintain pluripotency when grown on a feeder layer consisting of senescent fibroblasts (Takahashi and Yamanaka, 2006). However, except for urodele amphibians (Walters and Yun, 2020), tetrapod vertebrates have poor regenerative ability and do not respond to a senescence signal as effectively as *Hydractinia*.

We suggest that senescence is an ancient mechanism, instructing cells adjacent to an injury site to prepare for a regenerative event. We also speculate that other consequences of senescence that have been observed in mammals, such as long-term retention and accumulation of senescent cells, aging, chronic inflammation, and cancer, are side effects that evolved later in the evolution of these lineages, perhaps as consequence of the increase in cell fate stability and morphological complexity. Understanding the senescent environment and its role in cellular plasticity could pave the way for new treatments to enhance regeneration in poorly regenerating mammals.

### Limitations of the study

Our study provides strong evidence for a role for senescence signaling in cellular reprograming in cnidarians. While evidence from the literature provides indications to the existence of similar phenomena in other animals, the degree to which components of the senescence signaling pathway are evolutionarily conserved across phyla is at present unclear. Furthermore, markers for cellular senescence are not universal and no definitive senescence marker has been identified. Finally, the existence of different senescence ‘types’ has been proposed (Varela-Eirin and Demaria, 2022). Future work on other metazoans will be required to establish the pan-animal role of senescence signaling.

## Acknowledgements

We thank all members of the Frank lab for discussions and advice. The NIH Intramural Sequencing Center (NISC) is kindly acknowledged for generating the RNA sequence data. Confocal images were taken at the Centre for Microscopy and Imaging Core Facility at NUI Galway. UF is a Wellcome Trust Investigator in Science (grant no. 210722/Z/18/Z), and work in the Frank lab is also funded by the NSF EDGE program (grant no. 1923259). MSS is a Human Frontier Science Program Long-Term Postdoctoral Fellow (grant no. LT000756/2020-L). GK is an Irish Research Council postdoctoral fellow (project ID GOIPD/2020/149). HRH is a doctoral student in the Science Foundation Ireland Centre for Research Training in Genomic Data Science (grant no. 18/CRT/6214). ADB is funded by the Intramural Research Program of the National Human Genome Research Institute, National Institutes of Health (project No. ZIA HG000140).

## Author contribution

MSS and UF conceptualized the study. MSS performed experiments. GK performed phylogeny and Tunel assays. HRH and F analyzed data. ADB supervised the generation of RNA-seq data and making these data publicly available as described below. MSS and UF designed the experiments, analyzed data, and wrote the paper. All authors commented on and approved the manuscript.

## Data availability

The *Hydractinia symbiolongicarpus* genome is available in the *Hydractinia* Genome Project Portal through the NIH National Human Genome Research Institute (https://research.nhgri.nih.gov/hydractinia/). Corresponding data are available in the NCBI Sequence Read Archive (SRA) under BioProject PRJNA807936. Data corresponding to RNA-seq has been deposited under BioProject PRJNA824817. Bioinformatics scripts are available at https://github.com/UriFrankLab/Hsym_Hypostome_DiffExpr

---

**Figure S1.**
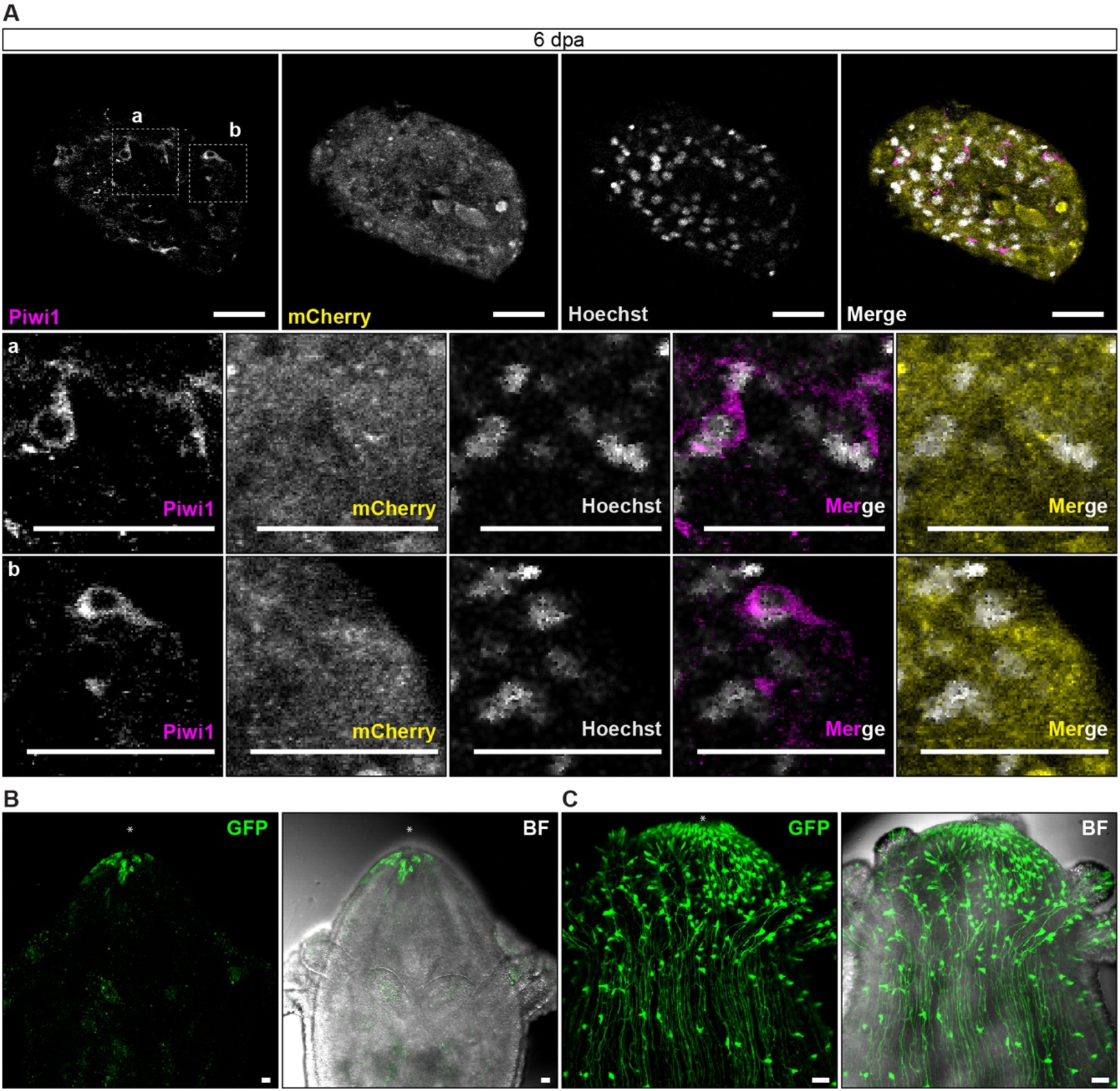
Data related to Figure 2. **(A)**. Single channel magnification of Figure 2D, showing the overlapping signal between mCherry and Piwi1 at 6dpa. **(B)** Wnt3:gfp intact polyp. **(C)** Rfamide:gfp intact polyp. Scale bars are 20 μm.

**Figure S2. Phylogenetic analyses of CDKN-like, mTor, and p53-like genes. (A-B)**, Topology of metazoan CDKN phylogeny obtained by Bayesian inference (A) and Maximum likelihood (B). Topologies show high divergence between CDKN sequences, numerous unresolved nodes, and inconsistency with species phylogeny. The three Hydractinia CDKI1, 2, and 3 named to indicate lack of clear orthologous relationships with vertebrate homologues. **(C-D)**, Topology of metazoan mTOR phylogeny obtained by Bayesian inference (C) and Maximum likelihood (D). The topologies are consistent with metazoans phylogeny: cnidarian sequences are monophyletics and the clade is sister-group to bilaterians. Inside bilaterians, two strongly supported monophyletic groups are identified, the deuterostomian and protostome genes (ecdysozoans+molluscs). Similarity between species phylogeny and mTOR phylogeny reflect the conservation of this gene during evolution. **(E-F)**, Topology of metazoan P53-like family phylogeny obtained by Bayesian inference (E) and Maximum likelihood (F). Bilaterian sequences are not monophyletic in Bayesian analysis, probably due to Long Branch attraction of ecdysozoan genes grouped with few cnidarian sequences. Most cnidarian sequences are distributed close to the root. In Maximum likelihood analysis, bilaterian genes are monophyletic and present coherent topology according to species phylogeny: deuterostomes are sister-group to protostomes (ecdysozoans+molluscs). P53, P63, and P73 are vertebrate specific paralogs. Consequently, the two *Hydractinia* P53 are named P53 like 1 and 2.

**Figure S3.**
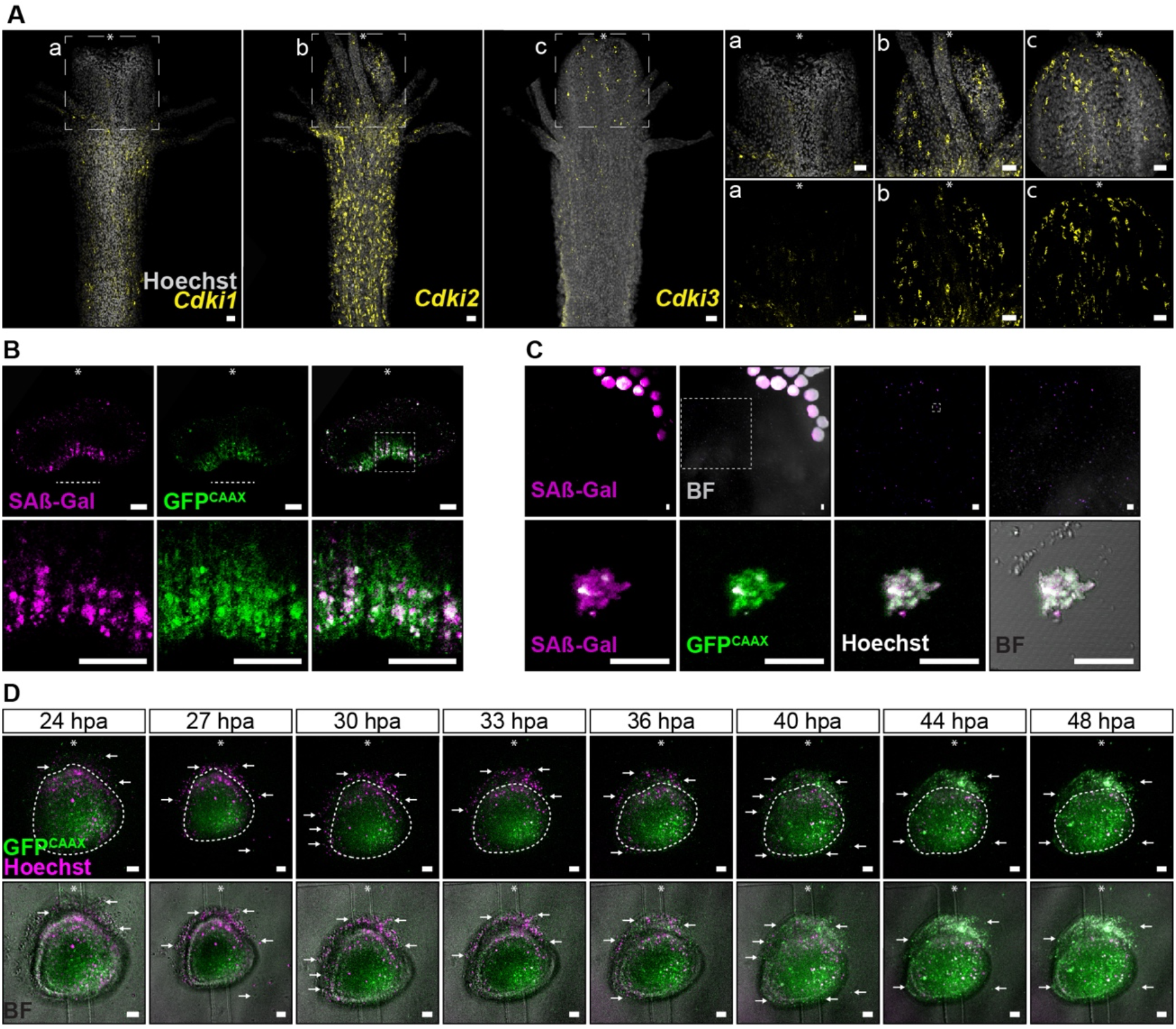
Expression patterns of *Cdki1-3* in intact polyps, and SA-βGal activity and expulsion of senescent cells during regeneration. **(A)** *Cdki1, Cdki2, and Cdki3* mRNA SABER-FISH in intact *Hydractinia* polyps. **(B)** SA-βGal and *Cdki1* are enhanced at the site of injury 1 dpa (dashed line). Asterisks indicate the oral pole pre-amputation. Lower panels correspond to magnifications of the region depicted with a dashed inset. **(C)** Cells expelled from regenerating hypostomes have high SA-βGal activity and and *Cdki1*::GFP^CAAX^espression. Lower panels correspond to magnifications of the region depicted with dashed insets. **(D)** Time-lapse imaging of an amputated *Cdki1*::GFP^CAAX^ reporter hypostome between 24 and 48 hpa. Arrows indicate expelled senescent cells from the tissue. Lower panel includes the brightfield channel. Time lapse of a second amputated *Cdki1*::GFP^CAAX^ reporter hypostome within 48 hpa. Arrows indicate the expel of senescent cells from the tissue. Scale bars: 20 μm

**Figure S4.**
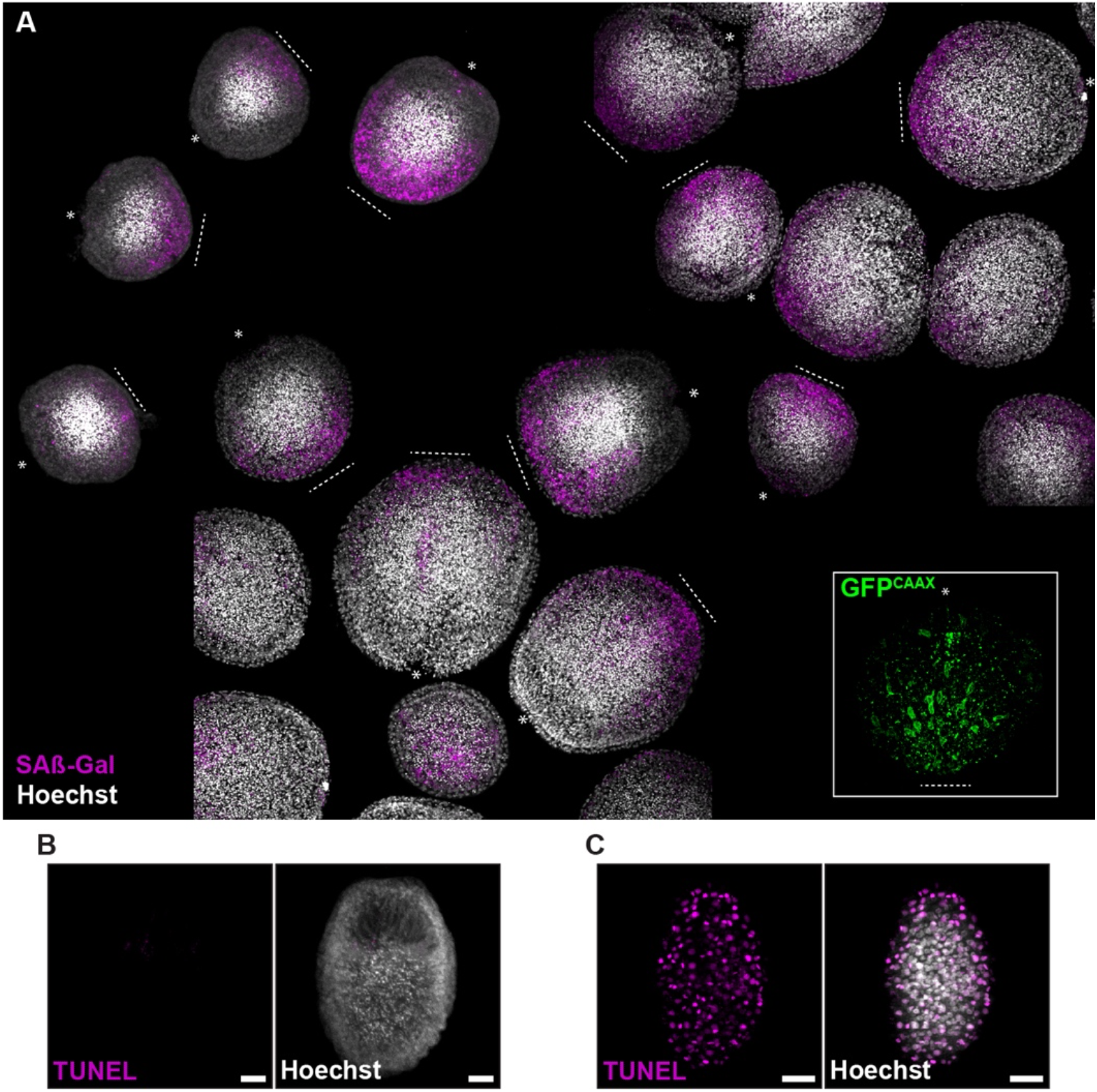
Cellular senescence markers are upregulated 1 dpa. **(A)** SA-βGal activity and *Cdki1* reporter GFP at the site of injury (dashed line). Asterisks indicate the oral pole pre-amputation. **(B)** Negative control for TUNEL staining. **(C)** Positive control for TUNEL staining

**Figure S5.**
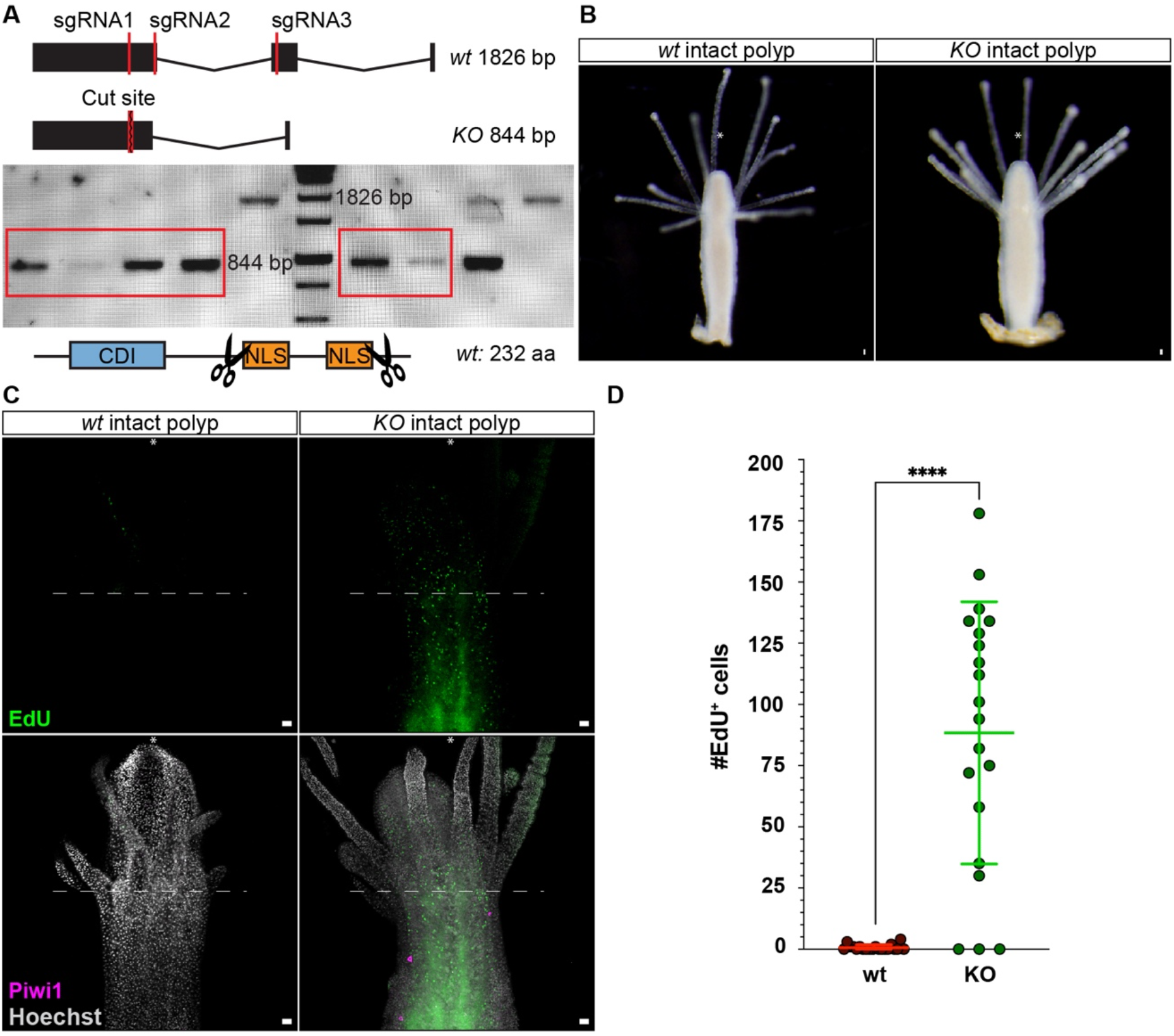
CRISPR-Cas9-mediated KO of th *Cdki1* gene. **(A)** The structure of *Cdki1* and the strategy used to knock it out. **(B)** Representative wild type and KO animals. **(C)** Distribution of S-phase and Piwi1^+^ cells in the upper body region of wild type and KO animals. **(D)** Quantification of the number of EdU^+^ cells observed in above the dashed line in C (p<0.0001, n=20). Scale bars: 20 μm

**Movie 1** – Time lapse of an amputated *Cdki1*::GFP^CAAX^ reporter hypostome corresponding to Figure 3F.

**Supplemental File 1** – DE genes between day 1 vs day 0, day 3 vs day 1, and day 6 vs day3.

**Supplemental File 2** – Quantification analyses and statistics.

## Methods

### *Hydractinia* husbandry

Adult *Hydractinia symbiolongicarpus* colonies were maintained as described in ref (Frank et al., 2020). Female and male colonies were grown on glass slides kept in artificial seawater (ASW) at 19-22ºC. Animals were fed four times per week with *Artemia* nauplii, and once a week with pureed oysters. To induce scheduled spawning, we kept the animals in a constant 14:10 light:dark cycle, where females and males spawn 1.5 hours after exposure to light.

### Hypostome isolation

One day starved adult *H. symbiolongicarpus* colonies were anaesthetized in 4% MgCl_2_ (in 50% distilled water / 50% filtered seawater). Polyps were dissected from the colony and decapitated. Hypostomes were isolated by removing the tentacles. Hypostomes were transferred to a glass Petri dish with freshly 0.2 μm filtered seawater and incubated at 22.4-23°C with constant reciprocal shaking in a temperature-controlled incubator. 0.2 μm filtered sea water was changed every two days.

### SpiDER-Gal staining

Hypostomes were collected and fixed in 0.2% glutaraldehyde (stock: 25%, Sigma-Aldrich; G5882), 4% paraformaldehyde (stock: 16%, Alfa Aesar; 43368) in filtered seawater for 2 minutes. Then, the fixative was replaced by 4% paraformaldehyde in filtered seawater and samples were incubated for 40 minutes. Samples were rinsed twice with seawater (3 minutes each) and proceed as per manufacturer’s indications. Samples were equilibrated with 1x Mcllvaine buffer, pH 6.0 (5x: 0.1 mol/l citric acid solution, 0.2 mol/l sodium phosphate solution) for 5 minutes. Then, samples were incubated with 1:500 SpiDER-Gal (1 mM; SG03-10, Dojindo Inc.) in 1x Mcllvaine buffer for 1 hour at RT. Samples were washed once with 1X PBST (0.3% TritonX-100 in 1X PBS) for 5 minutes, stained with Hoechst (20 mg/mL) 1:2000 in 1X PBST for 15 minutes, washed with 1X PBST, and mounted for imaging. Live animal staining with SpiDER-Gal was carried out in seawater.

### RNA Isolation from hypostome tissue

Total RNA was extracted from isolated hypostomes collected from wild type *H. symbiolongicarpus* strain 291-10. Samples consisted of three replicates of 220 hypostomes each at 0-, 1-, 3-, and 6-days post amputation (dpa). RNA was isolated using Direct-zol™ RNA MiniPrep kit (Zymo Research; R2050) per the manufacturer’s instructions. The eluted RNA was quantified using NanoDrop and the quality was checked using agarose-formaldehyde gel electrophoresis. After RNA isolation, samples were shipped to the NIH Intramural sequencing (NISC) for further processing and sequencing. RNA was amplified with the Ovation RNA-Seq System V2 kit, and sequencing libraries were made with Illumina TruSeq Stranded mRNA Library Prep Kit. Libraries were sequenced on one lane of Illumina NovaSeq 6000 (2×151 bp), generating between 87 and 137 million reads per sample (average 112 million reads). Only two replicates for 6 dpa were sequenced.

### Differential expression and gene set enrichment analysis

Raw reads were trimmed and aligned to the reference genome using Trim Galore https://www.bioinformatics.babraham.ac.uk/projects/trim_galore/ and HISAT2 (Kim et al., 2015), respectively. A count matrix of mapped reads per genomic feature was generated using Subread featureCounts (Liao et al., 2014) and converted into a DESeqDataSet. This dataset was used as an input to DESeq2 (Love et al., 2014) in order to generate a list of differentially expressed genes for each time point compared to the subsequent time point. Lists of differentially expressed genes were then separated based on up vs down regulated genes; sorted from the highest log2-fold changes to the lowest. These lists then used as an input for gene set enrichment analysis. To assign KEGG_Pathway terms to each gene ID, we fed *Hydractinia symbiolongicarpus* protein sequences, as coded in the recent genome (https://research.nhgri.nih.gov/hydractinia/download/protein_models/symbio/Hsym_primary_v1.0.aa.gz), into EggNOG-Mapper v5.0 (Huerta-Cepas et al., 2019), which then used as reference in ClusterProfiler. Gene set enrichment analysis were performed by using lists of differentially expressed genes as input into ClusterProfiler::enricher (Wu et al., 2021). We then plotted the identified enriched pathways (adj.p-value < 0.05) in Microsoft Excel.

### Transgenic animals

The generation of stable transgenic reporter animals was carried out as previously described (Künzel et al., 2010). Wnt3::GFP (DuBuc et al., 2020) and RFamide::GFP (Chrysostomou et al., 2022) reporters were previously generated in our lab. We used the GENEius online tool from Eurofins Genomics to codon optimize all the synthetic gene sequences for *Hydractinia symbiolongicarpus*. All synthetic genes were synthetized using IDT gBlocks gene fragments. Vector maps can be found below.

Fast Fluorescent timer protein reporter (FastFT): The codon optimized FastFT coding sequence (Subach et al., 2009) was designed in frame with P2A peptide and membrane GFP sequences (FastFT-P2A-GFP^CAAX^). The synthetic sequence was amplified by PCR and inserted into the Piwi1 reporter plasmid (Bradshaw et al., 2015) using *NotI* and *SacI* restriction enzymes. G0 colonies were bred to G2 (non-mosaic transgenic offspring). The timer property of the FastFT fluorescent protein (earlier blue and later red fluorescent forms according to maturation state) was aimed to differentiate i-cells (recently translated protein) from their progeny. The mature red form was detected using an mCherry filter set. However, our microscopes lacked the proper filter setup to detect the FastFT blue form. Thus, to identify i-cells, we calibrated the laser power to higher GFP intensities (DuBuc *et al*., 2020) that were devoid of or low in mCherry fluorescence (FastFT mature red form). Cdki1 reporter: 5’ upstream and 3’ downstream regulatory sequences of the *Cdki1* gene (*HyS0010*.*253*) were cloned from extracted genomic DNA by PCR and inserted into an open cloning vector. Membrane GFP sequence (GFP^CAAX^) was placed in frame with the 5’ upstream regulatory sequence. G0 colonies were bred to G2 (non-mosaic transgenic offspring).

OptoSOS: The OptoSos coding sequence (Johnson et al., 2017) was codon optimized and designed replacing the RFP sequence by our codon optimized mScarlet fluorescent protein (DuBuc *et al*., 2020). The synthetic sequence was amplified by PCR and inserted into the ß-tubulin reporter construct (DuBuc *et al*., 2020) using *NotI* and *SacI* restriction enzymes. We used BLAST to identify the endogenous *H. symbiolongicarpus* SOS catalytic domain that was amplified by PCR and inserted in frame with the SSPB sequence. Single injected embryos were grown to adult mosaic colonies, and we selected the animals who expressed the transgene in the hypostome.

### CRISPR/Cas9 knock-out

Single guide RNAs (sgRNAs) were designed using Geneious (2017.9.1.8) and CRISPRscan (Moreno-Mateos et al., 2015) tools. The three selected sgRNAs targeting *Cdki1* did not match other genomic sequences. Modified synthetic sgRNAs were synthesized by Synthego Inc and diluted according to the manufacturer recommendations. To form the RNP complex with Cas9, the three sgRNAs were incubated together (500 ng/μl total) with recombinant Cas9 (1μg/μl ; IDT, Cat. #1074181) for fifteen minutes at RT prior to being microinjected into zygotes.

### Single polyp genomic DNA extraction

Protocol was adapted from ref (DuBuc *et al*., 2020). A single polyp was isolated from a colony and transferred to 1.5 ml Eppendorf tube. The polyp was suspended in 50 μL Lysis buffer (10 mM Tris pH 8.0, 10 mM EDTA, 2% SDS), flicked few times, supplemented with 50 μL of digestion buffer (10% SDS, 0.4 mg/mL Proteinase K), and incubated at 56°C for 2-3 hours with occasional flicking. Then, 100 μL of cold phenol-chloroform (pH 8.0) was added. Samples were centrifuged at maximum speed for 15 minutes at 4°C, and supernatant was transferred to a new tube. Genomic DNA precipitation was carried out overnight at -20°C by adding two volumes of isopropanol and 1:10 total volume of Sodium Acetate Solution (3 M). Samples were cleaned up at 4°C by centrifugation and a 70% Ethanol wash. Extracted genomic DNA was dried out, resuspended in 10 μL nuclease-free water, and stored for further genotyping PCR.

### Genotyping CRISPR/Cas9 knock-outs

One hundred CRISPR-Cas9 larvae that had been injected as zygotes were metamorphosed and grown into small colonies. Of these, forty animals were analyzed for *Cdki1* mutations. Primers spanning the entire coding region of the gene were used in a PCR reaction to identify large deletions. We kept and grew eight G0 mosaic colonies to sexual maturity and crossed one male with a wild-type female. One hundred G1 animals were metamorphosed, grown to small colonies, and genotyped. Two sets of primers were used to identify heterozygous colonies carrying the genomic deletion: one spanning the entire coding region and another internal primer targeting the deleted intron. PCR products of heterozygous colonies were cloned into PGEMT-easy vector and sequenced using the T7 and SP6 primers. Each of these animals was a heterozygote carrying a 3’ deletion mutation in one *Cdki1* allele at the predicted cut sites and one wild type allele. One male and one female were crossed and G2 offspring were genotyped identifying five mutant homozygous animals. We performed genotyping on these animals from two independent DNA extractions and confirmed the mutation by sequencing.

### SABER-FISH

Oligo probes were prepared according to (Kishi et al., 2019) using hairpins 27 and 30. Tissue was fixed and dehydrated as previously described in (Chrysostomou *et al*., 2022; DuBuc *et al*., 2020): Tissue samples were incubated for 15 minutes in 4% MgCl2 (in 50% distilled water / 50% filtered seawater) and fixed in two steps. Fixative one: ice cold 0.2% glutaraldehyde (stock: 25%, Sigma-Aldrich; G5882), 4% paraformaldehyde (stock: 16%, Alfa Aesar; 43368) in filtered seawater for 90 seconds. Fixative two: 4% paraformaldehyde in in PBS-0.1% Tween (PTW) for one hour at 4°C. After three quick washes with PTW, samples were dehydrated in increasing concentrations of methanol in PTW and processed immediately by incubating in 1% H_2_O_2_ diluted in 100% MeOH (ice cold) for 45 min at 4°C. Followed by 2 quick washes with ice-cold 100% MeOH, samples were permeabilized in 75% acetone in 25% methanol and rehydrated into PTW. Two washes with glycine in PTW (2 mg/ml, 5min each; Fisher Scientific; BP381-1) followed by three PTW washes at RT. Samples were then washed three times with 1% (vol/vol) Triethanolamine pH 8.0 (TEA) in PTW for 5 min. 6 μl and 12 μl of acetic anhydride were added to the second and third TEA washes, respectively. Samples were then rinsed with PTW and incubated in pre-warmed Whyb buffer (2×SSC pH 7.0, 1% Tween-20, 40% Formamide) at 43°C for 10 min. Pre-hybridization and following steps were performed using pre-warmed reagents at 43°C according to (Kishi *et al*., 2019). Whyb buffer was replaced with Hyb1 buffer (2×SSC pH 7.0, 1% Tween-20, 40% Formamide, 10% Dextran sulfate) and incubated at 43°C overnight (pre-hybridization). Samples were transfer to Hybe buffer (40% Formamide, 5x SSC, 0.05mg/ml Heparin, 0.25% Tween20, 1% SDS, 1 mg/ml Salmon Sperm DNA, 1 mg/ml Roche blocking buffer powder) containing oligo probes at a concentration of 1μg/120μl and hybridized for two days at 43°C. Probes were removed and replaced with Whyb buffer for 10 min, followed by two washes with Whyb buffer (30 min each), a 10 min wash with 50% Whyb buffer in 2x SSCTw (2×SSC pH 7.0, 0.1% Tween-20), and two washes with 2X SSCTw (10 min each). Then, 2x SSCTw was replaced with PTW (two quick washes). Samples were warmed up to 37°C, prewarmed Hyb2/fluor solution (1×PBS, 0.2% Tween-20, 10% Dextran sulfate/10 μM Fluor Oligo) was added and incubated for one hour at 37°C. Hyb2/fluor solution was replaced with pre-warmed Whyb2 (1×PBS, 0.1% Tween-20, 30% Formamide) for a 10 min incubation at 37°C followed by two washes of 5 min each with PTW. Nuclear staining was then performed at RT by diluting Hoechst in PTW (1:2000) and incubating the samples for 45-60 min at RT. Samples were then quickly washed twice with PTW and mounted in 97% TDE. Samples were imaged within 4 days.

### Optogenetics

Isolated hypostomes from OptoSOS transgenic animals were exposed to 12 hours of constant blue illumination using ReefLed™ 50 (RedSea) lamp. Exposure time, light intensities, and distance from the light source (6 cm) was determined experimentally to optimal survival rate. Higher intensity values and longer exposure times resulted in lethality. Two control experiments were carried out alongside the light induction: (1) OptoSOS isolated hypostomes under dark conditions, and (2) wild type hypostomes exposed to blue light. After the light induction period, samples were incubated without blue light until day 6 post amputation and fixed for immunofluorescence (IF).

### Drug treatments

We incubated isolated hypostomes in 1 μM of Rapamycin (Sigma-Aldrich; 553210) or 1 μM of Navitoclax ABT-263 (MedChemExpress; HY-10087) for 28 hours. Controls were incubated in equivalent concentrations of DMSO. Treatment time was determined experimentally to optimize survival rates. A portion of the samples was fix at 24 hours post amputation for SpiDER-Gal staining. After treatment, samples were rinsed with fresh seawater, incubated until day 6 post amputation, and fixed for IF.

### Cellular staining

Immunofluorescence (IF) staining was performed as previously described in (DuBuc *et al*., 2020): the tissue was fixed in 4% Formaldehyde, 10 μl/mL Acetic acid (glacial) in filtered seawater (FSW) for 90 minutes at RT, followed by three washes with 1X PBST (20 minutes each). PBST was replaced with 5% normal goat serum (NGS; diluted in PBST) and fixed samples were blocked for 1 to 2 hours at room temperature with gentle rocking. For γH2AX staining, the tissue was fixed in Fix-2 from (Salinas-Saavedra et al., 2018) (100 mM HEPES pH 6.9, 0.05M EGTA, 5 mM MgSO4, 200 mM NaCl, 1x PBS, 3.7% Formaldehyde, 0.05% Glutaraldehyde, 0.5% Triton X-100, and FSW) followed by alcohol dehydration prior to PBST washes. Primary antibodies were diluted in 5% NGS to desired concentration (anti-Piwi1 1:500; anti-mCherry 1:100). Blocking solution was removed, replaced with primary antibodies diluted in NGS, and incubated overnight at 4°C. Samples were washed three times with PBST (10 minutes each), secondary antibodies were then applied (1:250 in 5% NGS) and incubated for 2 hours at RT. Tissue was rinsed with PBST and stained with Hoechst (20 mg/mL) 1:2000 in 1X PBST for 15 minutes, washed with 1X PBST, and mounted for imaging.

### EdU staining

EdU staining was done as previously described in (Bradshaw *et al*., 2015). Isolated polyps and hypostomes were incubated in 0.01 mM EdU in FSW for 30 minutes, rinsed with FSW and incubated in MgCl_2_ before fixation. Hypostomes were incubated in EdU for 2 days (3-4 dpa). Samples were fixed in 4% Formaldehyde, 10 μl/mL Acetic acid (glacial) in FSW for 90 minutes at room temperature (RT) and washed once with 3% BSA in PBST for 30min. Next, two washes with PBST for one hour and 30 minutes, respectively, followed by two washes with 3% BSA in PBST (5 minutes each). The tissue was then incubated in Click-iT cocktail for 30 minutes, followed by three washes of 3% BSA in PBST (20 minutes each).

### TUNEL labelling

TUNEL assays were performed using TM red In Situ Cell Death Detection Kit (#12156792910, Roche). Samples were fixed in 4% PFA in FSW for 90 minutes, then washed three times with PBS 0.01%Triton for 15 minutes, permeabilised with PBS 0.5%Triton for 90 minutes, and washed three times with 3% BSA in PBS for 15 minutes. Samples were incubated for 45 minutes at 37°C in a 50μL mix composed of 25 μL of reaction mix (Enzyme solution plus Label solution) and 25μL of 3% BSA in PBS. Negative controls were incubated in Label solution only and positive controls were incubated for 25 minutes at 37°C in DNAse I solution (Thermo Fisher Scientific, #EN0521) prior to incubation in Label solution only. After the reaction, samples were washed in PBS and nuclei stained using Hoechst.

### Imaging

Confocal images for IF and SABER-FISH were collected using an inverted Olympus Fluoview 1000 and Olympus Fluoview 3000 laser scanning confocal microscopes. For *in vivo* imaging, polyps from transgenic colonies were embedded in 0.8% low-melt agarose in filtered ASW and incubated in 35 mm imaging dishes with a glass bottom (Ibidi; D 263). Andor spinning disc and Olympus Fluoview 1000 laser scanning confocal microscopes were used to generate time-lapse movies and specific timepoint images. Raw images were visualized and analyzed using ImageJ/Fiji software and imaris viewer. Final figures were assembled using Adobe Illustrator and Adobe Photoshop.

*Piwi1* is expressed exclusively in i-cells. However, residual mRNA, protein, and the corresponding GFP reporter are present in decreasing concentrations in their progeny (DuBuc *et al*., 2020). Hence, we calibrated the laser power to higher *Piwi1*^+^ intensity signals using an intact polyp. We repeated this procedure prior to every acquisition to discriminate between i-cells (higher intensities) and their progeny (lower intensities).

### Cell count and statistical analyses

Stained cells of fixed hypostomes were counted with the Fiji cell counter tool using the raw source data. Statistical analyses were executed using GraphPad prism software. We plotted the cell numbers of the different conditions and differences were assessed by comparing medians using Mann-Whitney U test.

### Phylogenetic analysis

Multiple amino-acid alignments were generated with MAFFT 71 using default parameters. Gaps were removed using Gblocks 0.91b2. Final alignments were composed of 1022, 155, and 174 amino acids for mTOR alignment, P53 family alignment, and CDKI family alignment, respectively. We used PhyML 3.13 with maximum-likelihood method and 1000 bootstrap replicates. The best amino-acid evolution models to conduct analysis were determined, using MEGA114, to be the LG model for mTOR and P53 family alignment, and the JTT model for CDKI family alignment. Bayesian analyses were performed using MrBayes (v3.2.6) under mixed model. One fourth of the topologies were discarded (burn-in values), and the remaining ones were used to calculate the posterior probabilities. Bayesian analysis for mTOR, P53 family, and CDKI family ran for 200,000 generations with 5 randomly started simultaneous Markov chains (MC), 500,000 generations with 10 MC, and 3,000,000 with 15 MC, respectively. For all analyses, the first chain was a cold chain and the others were heated chains.

## Key resources table

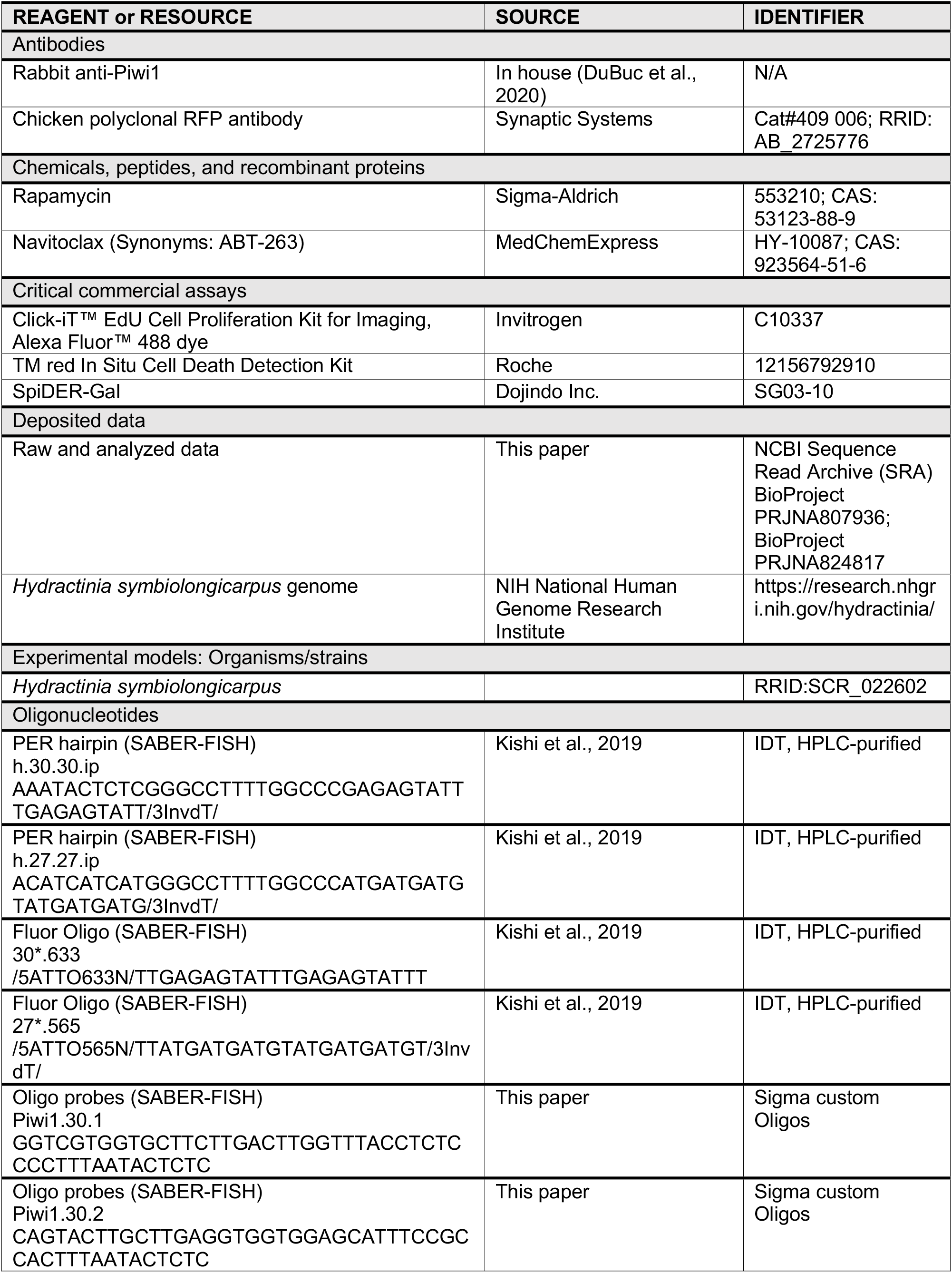
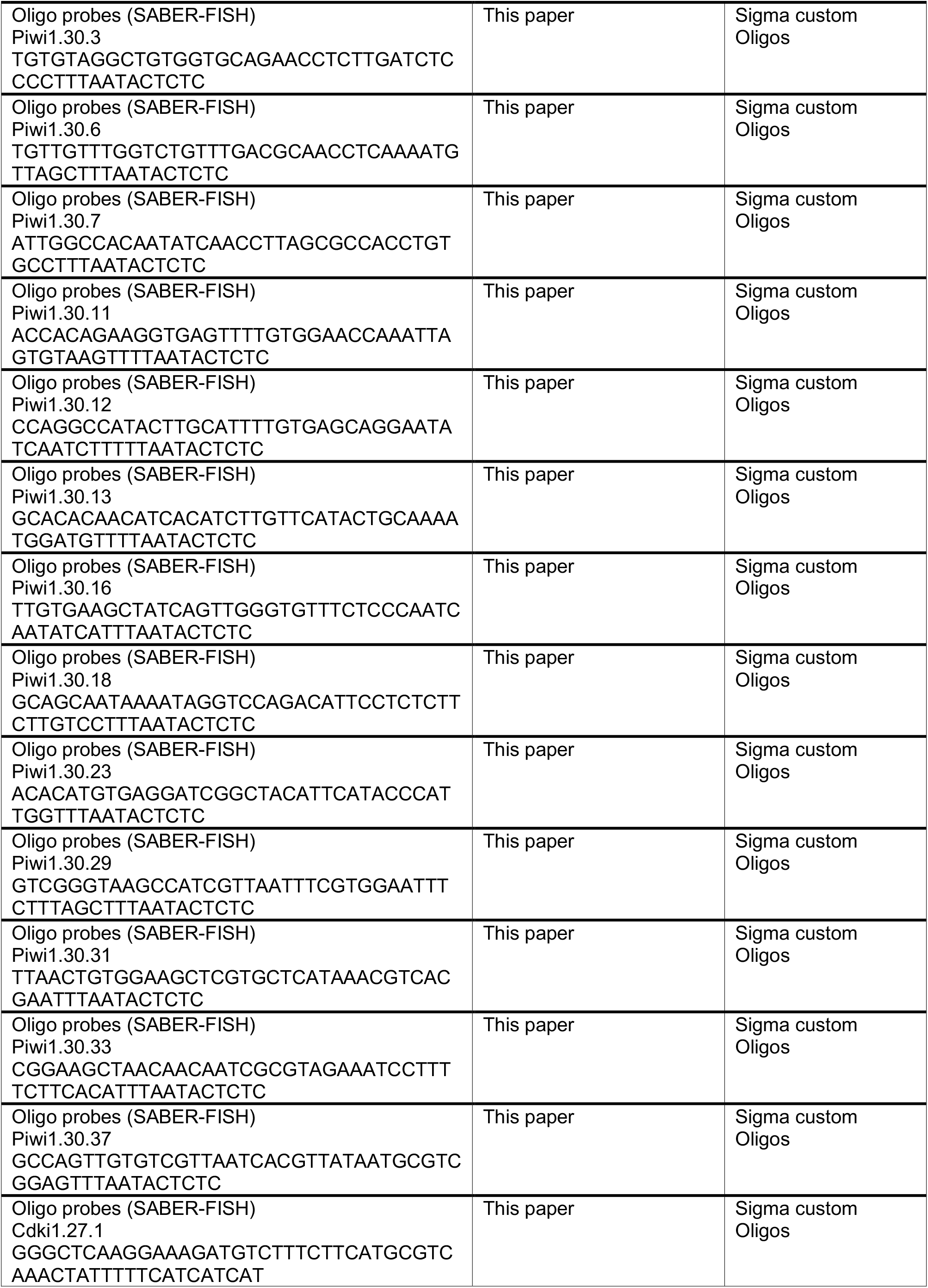
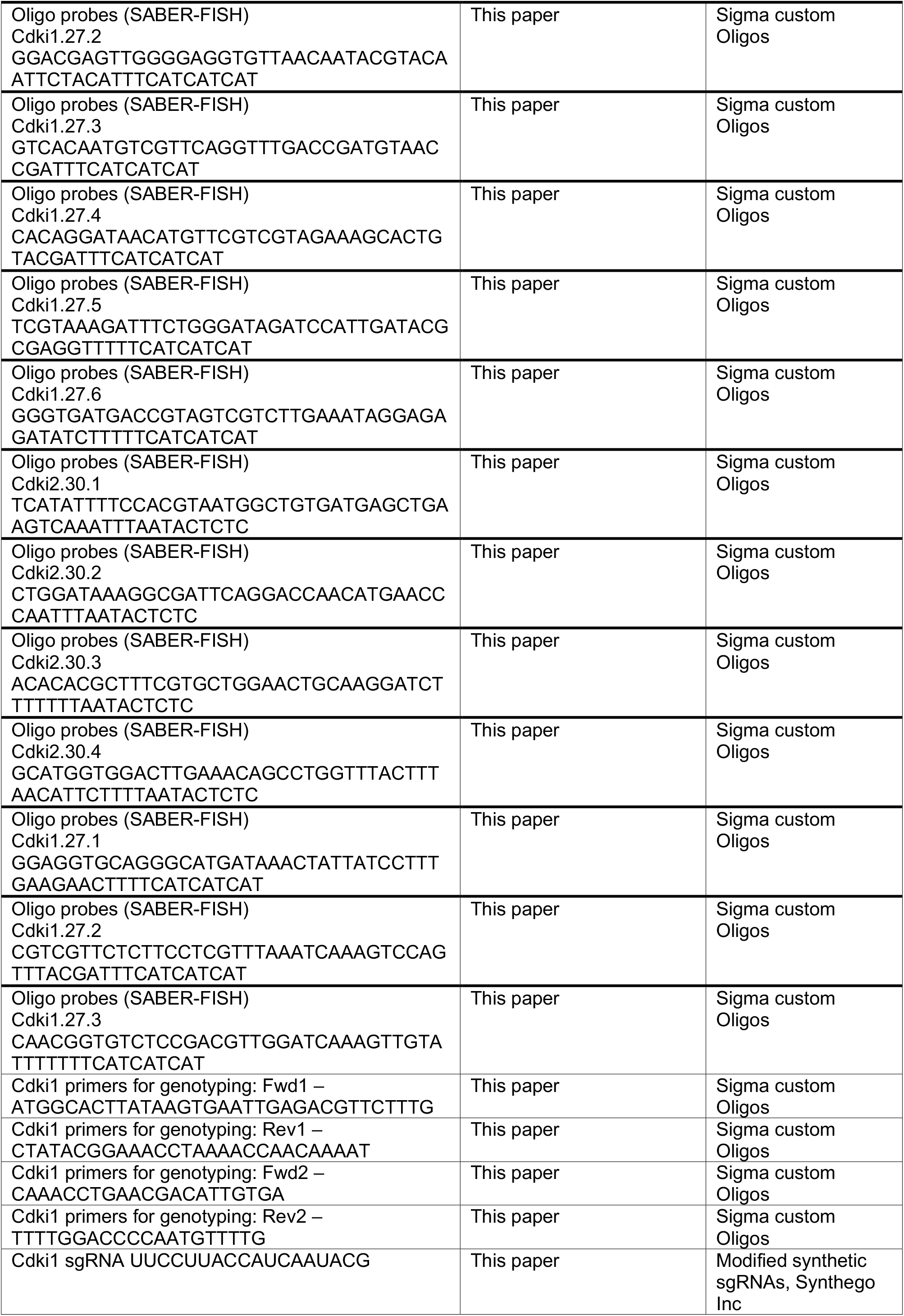
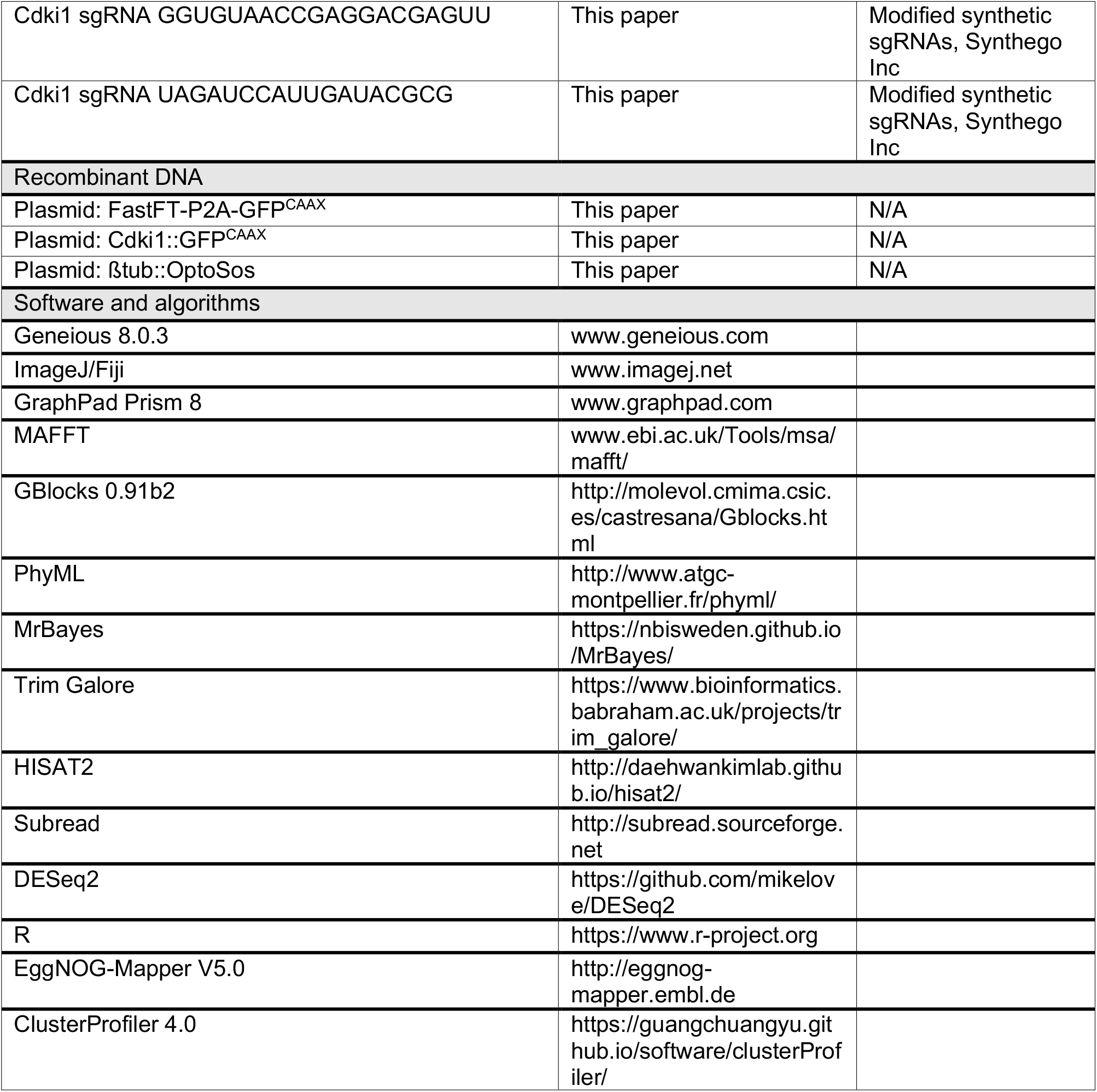

## Notes

### Competing Interest Statement

The authors have declared no competing interest.

